# Impact of transposable elements on genome structure and evolution in bread wheat

**DOI:** 10.1101/363192

**Authors:** Thomas Wicker, Heidrun Gundlach, Manuel Spannagl, Cristobal Uauy, Philippa Borrill, Ricardo H. Ramírez-González, Romain De Oliveira, International Wheat Genome Sequencing Consortium, Klaus F. X. Mayer, Etienne Paux, Frédéric Choulet

## Abstract

**Background:** Transposable elements (TEs) are ubiquitous components of genomes and they are the main contributors to genome evolution. The reference sequence of the hexaploid bread wheat (*Triticum aestivum* L.) genome enabled for the first time a comprehensive genome-wide view of the dynamics of TEs that have massively proliferated in the A, B, and D subgenomes.

**Results:** TEs represent 85% of the genome. We traced back TE amplification dynamics in the evolutionary history of wheat and did not find large bursts in the wake of either the tetra- or the hexaploidization. Despite the massive turnover of TEs since A, B, and D diverged, 76% of TE families are present in similar proportions in the three subgenomes. Moreover, spacing between homeologous genes was also conserved. TE content around genes is very different from the TE space comprising large intergenic regions and families that are enriched or depleted from gene promoters are the same in the three subgenomes.

**Conclusions:** The chromosome-scale assembly of the wheat genome provided an unprecedented genome-wide view of the organization and impact of TEs in such a complex genome. Our results suggest that TEs play a role at the structural level and that the overall chromatin structure is likely under selection pressure.

## Background

Transposable elements (TEs) are ubiquitous components of genomes and one of the major forces driving genome evolution [1]. They are classified into two classes: retrotransposons (class 1), transposing via reverse transcription of their mRNA, and DNA transposons (class 2), representing all other types of elements [2]. TEs are small genetic units with the ability to make copies of themselves or move around in the genome. They do not encode a function that would allow them to be maintained by selection across generations rather, their strategy relies on their autonomous or non-autonomous amplification. TEs are subject to rapid turn-over, are the main contributors of intraspecific genomic diversity, and are the main factor explaining genome size variations. Thus, TEs represent the dynamic reservoir of the genomes. They are epigenetically silenced [3], preventing them from long-term massive amplification that could be detrimental. The dynamics of TEs in genomes remains unclear and it was supposed that they may escape silencing and experience bursts of amplification followed by rapid silencing. Their impact on gene expression has also been documented in many species (for review [4]). In addition, they play a role at the structural level, as essential components of centromeric chromatin in plants [3, 5]. Plant genomes are generally dominated by a small number of highly repeated families, especially class I Gypsy and Copia LTR-retrotransposons [6-10]. Most of our knowledge about TE dynamics and their impact on gene expression in complex plant genomes comes from maize [10-14]. At the whole genome level, Makarevitch et al. have shown that four to nine maize TE families, including all major class I superfamilies (Gypsy, Copia, LINEs) and DNA transposons, are enriched (>2 fold) in promoters of genes being up-regulated in response to different abiotic stresses [15]. This study also suggested that TEs are a major source of allelic variations explaining differential response to stress between accessions.

The genome of bread wheat (*Triticum aestivum* L.), one of the most important crop species, has also undergone massive TE amplification with over 85% of it being derived from such repeat elements. It is an allohexaploid comprised of three subgenomes (termed A, B, and D) that have diverged from a common ancestor around 2-3 million years ago (according to molecular dating of chloroplast DNA [16]) and hybridized within the last half million years. This led to the formation of a complex, redundant, and allohexaploid genome. These characteristics make the wheat genome by far the largest and most complex genome that has been sequenced and assembled into near complete chromosomes so far. They, however, also make wheat a unique system to study the impact of TE activity on genome structure, function and organization.

Previously only one reference sequence quality wheat chromosome was available which we annotated using our automated TE annotation pipeline (CLARITE) [17, 18]. However, it was unknown whether the TE content of chromosome 3B was typical of all wheat chromosomes and how TE content varied between the A, B, and D subgenomes. Therefore, in this study, we address the contribution of TEs to wheat genome evolution at a chromosome-wide scale. We report on the comparison of the three A-B-D subgenomes in terms of TE content and proliferation dynamics. We show that, although TEs have been completely reshuffled since A-B-D diverged, their proportions are quite conserved between subgenomes. In addition, the TE landscape in the direct vicinity of genes is very similar between the three subgenomes. Our results strongly suggest that TEs play a role at the structural level, and that the overall chromatin structure is likely under selection pressure. We also identified TE families that are overrepresented in promoters compared to the rest of the genome, but did not reveal a strong association between particular TE families and nearby gene expression pattern, nor with stress-response.

## Results and discussion

### TE content and distribution along the 21 bread wheat chromosomes

Building from a decade-long effort from the wheat genomics community, we used the accumulated knowledge about TEs to precisely delineate the TE repertoire of the 21 chromosomes based on similarity search with a high quality TE databank: ClariTeRep [17] which includes TREP [19]. CLARITE was used to model TEs in the sequence and their nested insertions when possible [17]. This led to the identification of 3,968,974 TE copies, belonging to 505 families, and representing 85% of RefSeq_v1.0. Overall, the TE proportion is very similar in the A, B, and D subgenomes as they represented 86%, 85%, and 83% of the sequence, respectively. However, the sizes of the subgenomes differ: with 5.18 Gb, the B subgenome has the largest assembly size, followed by the A subgenome (4.93 Gb) and the smaller D subgenome (3.95 Gb). The repetitive fraction is mostly dominated by TEs of the class I Gypsy and Copia and class II CACTA superfamilies, while other superfamilies contribute only very little to overall genome size (Table 1, Fig. 1A).

**Figure 1:**
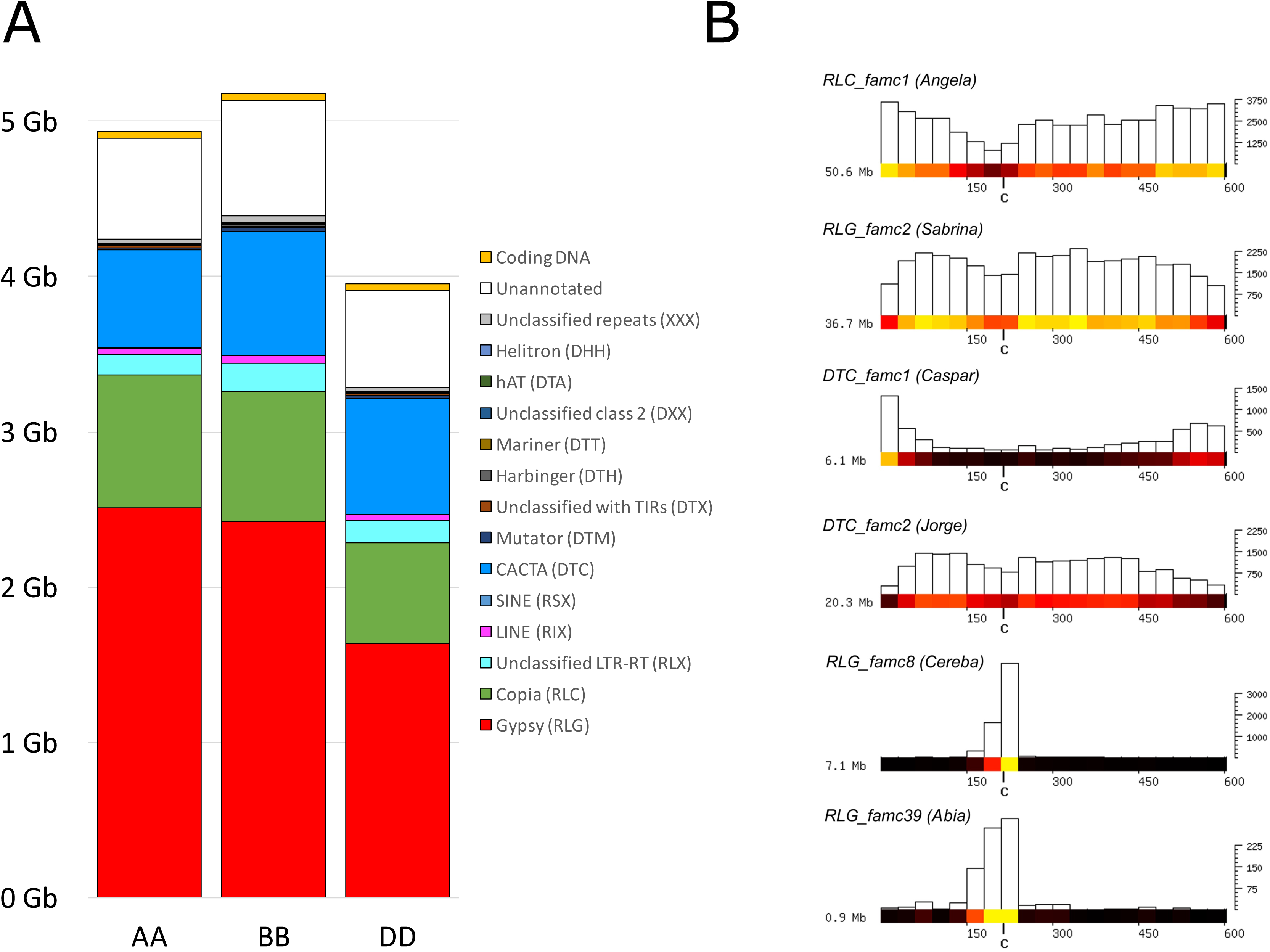
TE composition of the three wheat subgenomes and examples of chromosomal distributions. **A.** Stacked histograms representing the contribution of each TE superfamily to the three subgenomes. Un-annotated sequences are depicted in white and coding exons (accounting only the representative transcript per gene) in orange. **B.** Distribution of TE subfamilies along wheat chromosome 1A (as a representative of all chromosomes). The full datasets are shown in Fig. S1-11. The TE distribution is shown in 30 Mb windows along chromosomes. TE abundance per 30 Mb-window is shown as a heat map and as a bar plot. The x-axis indicates the physical position in Mb, while the y-axis indicates the number of kb the TE family contributes to each 30 Mb. The total contribution in Mb of the respective TE family to the chromosome is depicted at the left.

**Table 1:** Proportion of TE superfamilies in the A, B, and D subgenomes and at the whole genome level. Proportions are expressed as the percentage of sequences assigned to each superfamily relatively to the genome size. TIR: terminal inverted repeat.

|  | A | B | D | Complete genome |
| --- | --- | --- | --- | --- |
| Class 1 LTR-retrotransposons |  |  |  |  |
| Gypsy (RLG) | 50.9% | 46.8% | 41.4% | 46.7% |
| Copia (RLC) | 17.5% | 16.2% | 16.3% | 16.7% |
| Unclassified LTR-RT (RLX) | 2.6% | 3.5% | 3.7% | 3.2% |
| Non-LTR-retrotransposons |  |  |  |  |
| LINE (RIX) | 0.82% | 0.96% | 0.93% | 0.90% |
| SINE (RSX) | 0.01% | 0.01% | 0.01% | 0.01% |
| Class 2 DNA transposons |  |  |  |  |
| CACTA (DTC) | 12.8% | 15.5% | 19.0% | 15.5% |
| Mutator (DTM) | 0.30% | 0.38% | 0.48% | 0.38% |
| Unclassified with TIRs (DTX) | 0.21% | 0.20% | 0.22% | 0.21% |
| Harbinger (DTH) | 0.15% | 0.16% | 0.18% | 0.16% |
| Mariner (DTT) | 0.14% | 0.16% | 0.17% | 0.16% |
| Unclassified class 2 (DXX) | 0.05% | 0.08% | 0.05% | 0.06% |
| hAT (DTA) | 0.01% | 0.01% | 0.01% | 0.01% |
| Helitrons (DHH) | 0.00% | 0.00% | 0.00% | 0.00% |
| Unclassified repeats (XXX) | 0.55% | 0.85% | 0.63% | 0.68% |
| Genes and non TE-related DNA | 13.9% | 15.3% | 16.8% | 15.2% |

At the superfamily level, the A, B, and D subgenomes have very similar TE compositions (Fig. 1A). Almost half of the size difference between A and B (106/245 Mb) is due to low-copy DNA rather than TEs. Since the amount of coding DNA is very conserved (43, 46 and 44 Mb, respectively), the difference is mainly due to parts of the genome that remained un-annotated so far. This un-annotated portion of the genome may contain highly degenerated TE copies or as yet uncharacterized TE types. The amount of DNA represented by each superfamily is also similar between A, B, and D subgenomes, except for Gypsy retrotransposons, which are less abundant in the D subgenome (1.6 Gb, compared to 2.5 and 2.4 Gb in A and B, respectively). This feature alone can explain why the D subgenome is ~900 Mb smaller. Indeed, 10 of the 14 Gypsy families that account for more than 100 Mb each are more abundant in the A and B than in D genomes, representing almost all of the subgenome size difference.

Similar to other complex genomes, only six highly abundant TE families represent more than half of the TE content: RLC_famc1 (Angela), DTC_famc2 (Jorge), RLG_famc2 (Sabrina), RLG_famc1 (Fatima), RLG_famc7 (Sumana/Sumaya), and RLG_famc5 (WHAM), while 486 families out of 505 (96%) account each for less than 1% of the TE fraction. In term of copy number, 50% (253) of the families are repeated in fewer than 1,000 copies at the whole genome level, while more than 100,000 copies were detected for each of the seven most repeated families (up to 420,639 *Jorge* copies).

Local variations of the TE density were observed following a pattern common to all chromosomes: the TE proportion is lower (on average 73%) in the distal regions than in the proximal and interstitial regions (on average 89%). However, much stronger local variations were observed when distributions of individual TE families were studied. Figure 1B shows TE distributions using chromosome 1A as a representative example. Distributions for selected TE families on all chromosomes are shown in Figures S1-S11. The most abundant TE family, RLC_famc1 (Angela) was enriched toward telomeres and depleted in proximal regions. In contrast, highly abundant Gypsy retrotransposons RLG_famc2 (Sabrina, Fig. 1B) and RLG_famc5 (WHAM, not shown) were enriched in central parts of chromosome arms and less abundant in distal regions. CACTA TEs also showed a variety of distribution patterns. They can be grouped into distinct clades depending on their distribution pattern as suggested earlier based on chromosome 3B TE analyses [17]. Families of the Caspar clade [20] are highly enriched in telomeric regions, as is shown for the example of the DTC_famc1 (Caspar) whereas DTC_famc2 (Jorge) showed the opposite pattern (Fig. 1B).

Centromeres have a specific TE content. Previous studies on barley and wheat reported that the Gypsy family RLG_famc8.3 (Cereba) is enriched in centromeres [21, 22]. It was speculated that Cereba integrase can target centromere-specific heterochromatin due to the presence of a chromodomain that binds specifically to centromeric histones [23]. We found that wheat Cereba elements are concentrated in centromeric regions while absent from the rest of the genome (Fig. 1B, Fig. S8), and so are their closely related subfamilies RLG_famc8.1 and RLG_famc8.2 (Quinta). We identified new TE families that are also highly enriched in centromeres. The family RLG_famc39 (Abia) is a relative of Cereba, although there is very little sequence DNA conservation between the two. However, at the protein level, Cereba is its closest homolog. Abia and Cereba have an extremely similar distribution (Fig. 1B, Fig. S8-S9), competing for the centromeric niche. This is apparent on chr6A where Cereba is more abundant while on 3B, Abia is more abundant. Abia seems to be a wheat specific TE family as it was not present in the recently published barley genome [24]. A recent study on the barley genome reported on a novel centromeric Gypsy family called Abiba [20]. We identified a homolog in wheat: RLG_famc40 (Abiba), with two distinct subfamilies RLG_famc40.1 and RLG_famc40.2, corresponding to the putatively autonomous and non-autonomous variants. Abiba is enriched in central parts of chromosomes but with a broader spreading compared to Abia and Cereba (Fig. S10-S11). At a higher resolution, we identified large tandem arrays of Cereba and Abia elements that correspond to the high kmer frequencies observed at the centromeres (Fig. 2D), which might be the signature of functional centromeres (Fig. S12).

**Figure 2:**
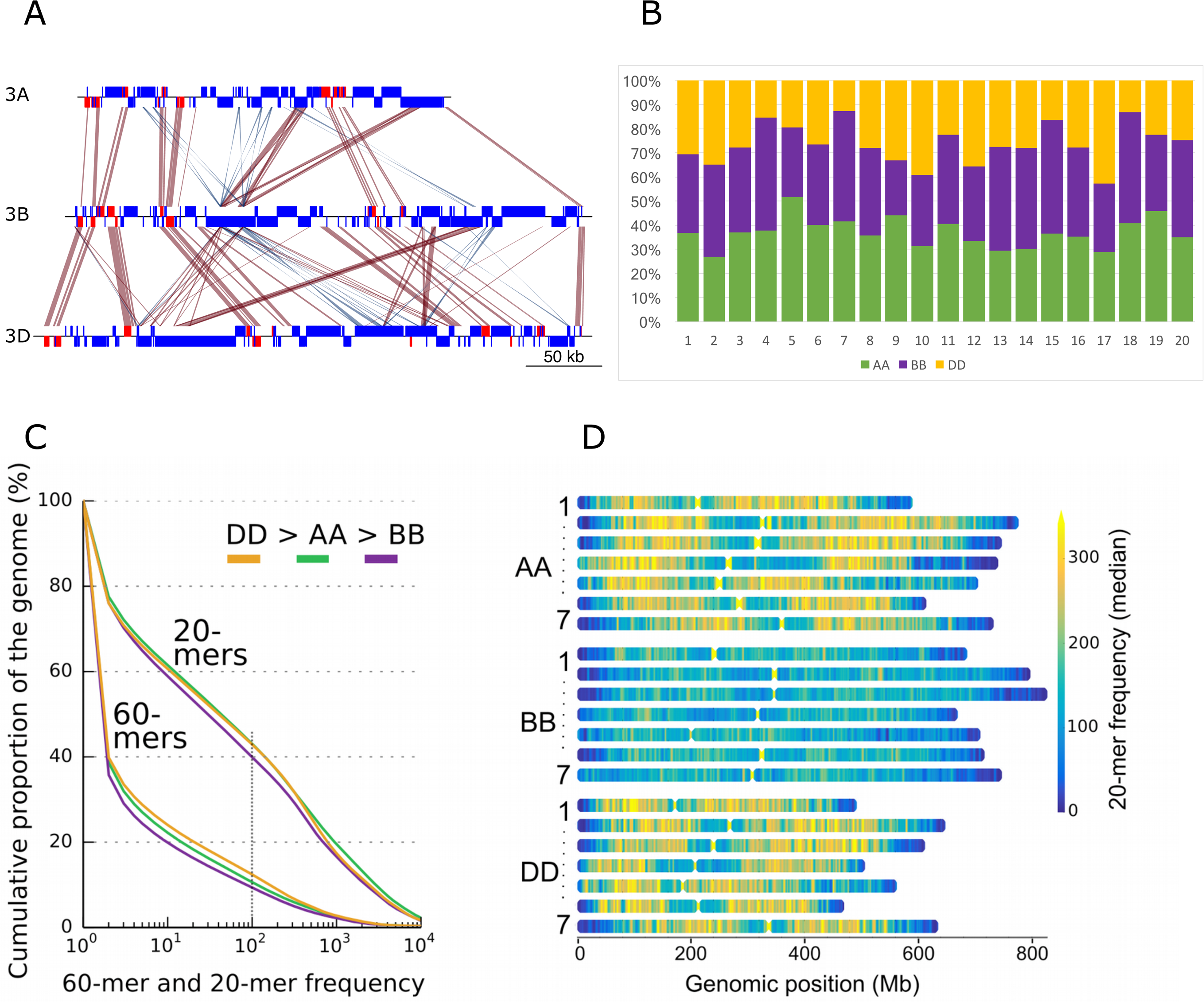
Variability and similarity of the repeat composition of the three wheat subgenomes. **A.** Example of sequence alignment of three homeologous regions of ca. 300 kb on chromosomes 3A (from 683.185 to 683.435 Mb), 3B (from 723.440 to 723.790 Mb), and 3D (from 546.330 to 546.700 Mb). Genes: red boxes; TEs: blue boxes. Sequences sharing >90% identity over more than 400 bp are represented by red (+/+ strand matches) and blue (+/− strand matches) areas. It shows the high conservation between homeologous genes and collinearity between A-B-D, and it shows the absence of conservation of the intergenic space while intergenic distances tend to be similar between homeologs. Similarities observed between TEs are not collinear and thus strongly suggest independent insertions, in the three subgenomes, of TEs from the same family instead of homeologous relationships. **B**. Proportions of the 20 most abundant TE families comprising the hexaploid wheat genome depicted as fractions of A, B, and D subgenomes. For each family, the A-B-D fractions are represented in green, violet, and orange, respectively. 1: RLC_famc1 (Angela WIS); 2: DTC_famc2 (Jorge); 3: RLG_famc2 (Sabrina Derami Egug); 4: RLG_famc1 (Fatima); 5: RLG_famc7 (Erika Sumana Sumaya); 6: RLG_famc5 (WHAM Wilma Sakura); 7: RLG_famc3 (Laura); 8: RLG_famc4 (Nusif); 9: RLG_famc11 (Romana Romani); 10: RLG_famc10 (Carmilla Ifis); 11: RLC_famc3 (Claudia Maximus); 12: RLG_famc13 (Latidu); 13: RLG_famc6 (Wilma); 14: RLG_famc9 (Daniela Danae Olivia); 15: RLC_famc2 (Barbara); 16: DTC_famc1 (Caspar Clifford Donald Heyjude); 17: RLG_famc14 (Lila); 18: RLG_famc15 (Jeli); 19: RLG_famc8 (Cereba Quinta); 20: DTC_famc6 (TAT1). **C.** Kmer-defined proportion of repeats of the subgenomes. Cumulative genome coverage of 20- and 60-mers at increasing frequencies. Around 40% of each subgenome assembly consist of 20-mers occurring >=100 times. At the 60-mer level the D subgenome has the highest and B the lowest proportion of repeats. **D**. Distribution of 20-mer frequencies across physical chromosomes. The B genome has the lowest overall proportion of repeats.

### Similarity and variability of the TE content between the A, B, and D subgenomes

A genome-wide comparative analysis of the 107,891 high confidence genes predicted along the A, B, and D subgenomes (35,345, 35,643, and 34,212, respectively) was described in detail in [25]. It revealed that 74% of the genes are homeologs, with the vast majority being syntenic. Thus, gene-based comparisons of A-B-D highlighted a strong conservation and collinearity between the three genomes. However, outside genes and their immediate surrounding regions, we found almost no sequence conservation of individual TE copies that shape the intergenic regions (Fig. 2A). This is due to “genomic turnover” [26] through rounds of TE amplifications followed by removal via unequal crossing-overs or deletions that occur during double-strand repair [27]. Thus, TEs have been massively reshuffled since the divergence of A, B, and D, raising the question of the extent of variability of the repeat fraction between subgenomes in wheat.

Analysis of kmer content of RefSeq_v1.0 showed that 20-mers occurring 100x or more cover around 40% of the wheat genome sequence (Fig 2C). Considering 60-mers, this value decreases to only 10%. This pattern was strongly similar between subgenomes, although a slight difference was observed: repeated kmers covered a larger proportion of the subgenome D>A>B. This lower proportion of repeats in the B subgenome is also obvious using a heat-map of 20-mer frequencies (Fig 2D), showing that the larger size of the B genome was due to a higher diversity of TE families.

We then compared A, B, and D subgenomes at the TE family level. We did not find any TE families (accounting >10 kb) that are specific for a single subgenome or completely absent in one. Only two cases of subgenome specific tandem repeats were found, XXX_famc46 and XXX_famc47. More surprisingly, the abundance of most TE families is similar in the A, B, and D subgenomes. Indeed, among the 165 families which represent at least 1 Mb of DNA each, 125 (76%) are present in similar proportions in the 3 subgenomes, i.e. we found less than a 2-fold change of the proportion between subgenomes. Fig. 2B represents the proportions of the 20 most abundant families in the 3 subgenomes which account for 84% of the whole TE fraction. Their proportion is close to the relative sizes of the three subgenomes: 35%-37%-28% for A-B-D, respectively. This highlighted the fact that, not only are the three subgenomes shaped by the same TE families, but also that these families are present in proportions that are conserved. Consistent with this, we identified only eleven (7%) TE families that show a strong difference (i.e. more than 3-fold change in abundance) between two subgenomes, representing only 2% of the overall TE fraction.

Thus, although TEs have been extensively reshuffled during the independent evolution of the diploid A-B-D lineages (Fig. 2A), and although TEs have transposed and proliferated very little since polyploidization (0.5 Mya, see below), the TE families that currently shape the three subgenomes are the same, and more strikingly, their abundance remains very similar. We conclude that almost all families ancestrally present in the A-B-D common ancestor have been active at some point and their amplification compensated their loss by deletion, thus suggesting a dynamic in which families are maintained at the equilibrium in the genome for millions of years. These findings disagree with the common notion that TEs evolve by successive amplification bursts of a limited number of active families before their silencing. For example, Piegu et al. showed that an amplification burst of a single retrotransposon family led to a near doubling of the genome size in *Oryza australiensis* [28]. In wheat, the high-quality reference genome assembly and detailed TE annotation revealed that the difference of size between subgenomes is not mainly explained by a few TE families that have experienced a massive and recent burst.

Instead, many TE families contribute to this phenomenon. The contribution of a wide diversity of families to the expansion of genomes was previously suggested for plants with very large genomes >30 Gb [29]. Our data show that this is the case also for the *Triticeae* tribe.

Although practically all TE families are found in similar abundance in the three subgenomes, subfamilies show strong differences in abundance in the A, B, and D genomes (Fig. 3). For example, the highly abundant RLC_famc1 (Fatima) family has diverged into at least five subfamilies (1.1 to 1.5). Only RLC_famc1.1 contains potentially functional reverse transcriptase (RT) and integrase (INT) genes while RLC_famc1.4 and RLC_famc1.5 contain gag and protease ORFs. RLC_famc1.2 and RLC_famc1.3 appear to be non-autonomous as they do not contain any intact ORFs. We suggest that RLC_famc1.1 provides functional RT and INT proteins while protease and GAG are provided by other subfamilies. Their contrasted abundance revealed that RLC_famc1.4 and RLC_famc1.5 proliferated specifically in the B and A lineages, respectively (Fig. 3A).

**Figure 3:**
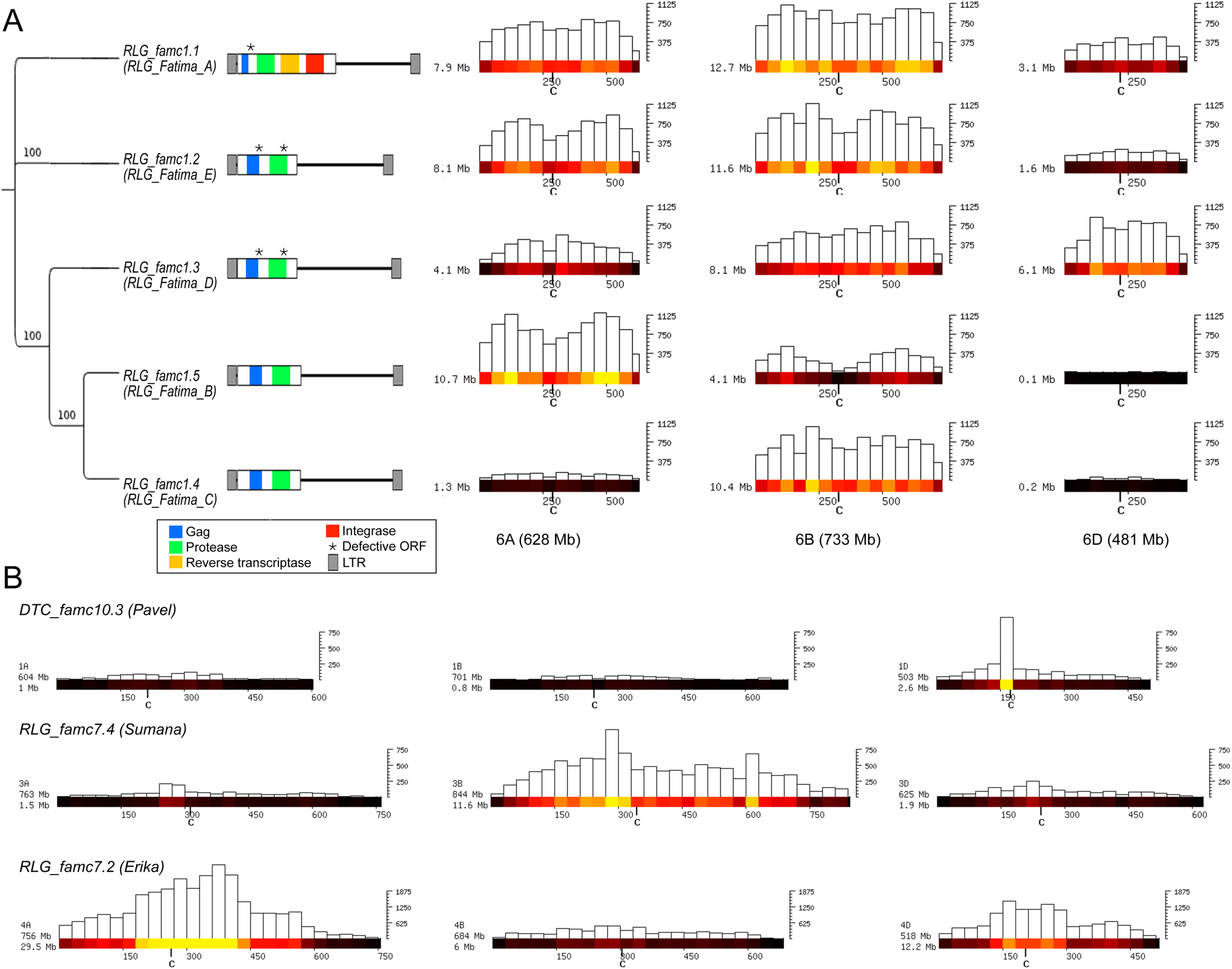
Distribution of different subfamilies in the A, B, and D subgenomes. **A**. Distribution of RLC_famc1 (Fatima) retrotransposons. Group 6 chromosomes were chosen as representative for the whole genome. A phylogenetic tree of the different subfamilies is shown at the left. For the construction of the phylogenetic tree, the LTR sequences were used (internal domains between RLC_famc1.1 and the other subfamilies are completely different, as only RLC_famc1.1 contains reverse transcriptase and integrase genes). Bootstrap values (100 repetitions) are indicated. Sequence organization and gene content of the individual subfamilies are shown to the right of the tree. Chromosomal distributions are shown at the right in bins of 50 Mb as heat maps and bar plots to indicate absolute numbers. The y-axis indicates the total number of kb that is occupied by the respective subfamily in each bin. The most recently diverged subfamilies RLC_famc1.4 and RLC_famc1.5 show strong differences in abundance in different subgenomes. **B**. Examples of TE subfamilies that have strongly differing copy numbers in the A, B, and D genomes. Again, only a single group of homeologous chromosomes is shown (see Fig. S1-3 for the other chromosomes). Abundance is shown in 30 Mb windows.

In total, we identified 18 different subfamilies (belonging to eleven different families) which show subgenome specific over- or under-representation (Table 2). Here, we only considered TE families that contribute more than 0.1% to the total genome and are at least 3-fold over- or under-represented in one of the subgenomes. This illustrated that these eleven highly abundant families did not show a bias between A-B-D at the family level, but are composed of several subfamilies that were differentially amplified in the three diploid lineages. The CACTA family DTC_famc10.3 (Pavel) is much more abundant in the D subgenome than in the A and B genomes (Fig. S1). Interestingly, the Pavel subfamily also seems to have evolved a preference for inserting close to centromeres in the D genome, while this tendency is not obvious in the A and B genomes (Fig. 3B). Generally, subfamilies were enriched in a single genome (Table 2). In only four cases, a subfamily was depleted in one subgenome while abundant at similar levels in the other two. Three of these cases were found in the D genome. This is consistent with the smaller D genome size and differences in highly abundant elements contribute to this difference.

**Table 2:**
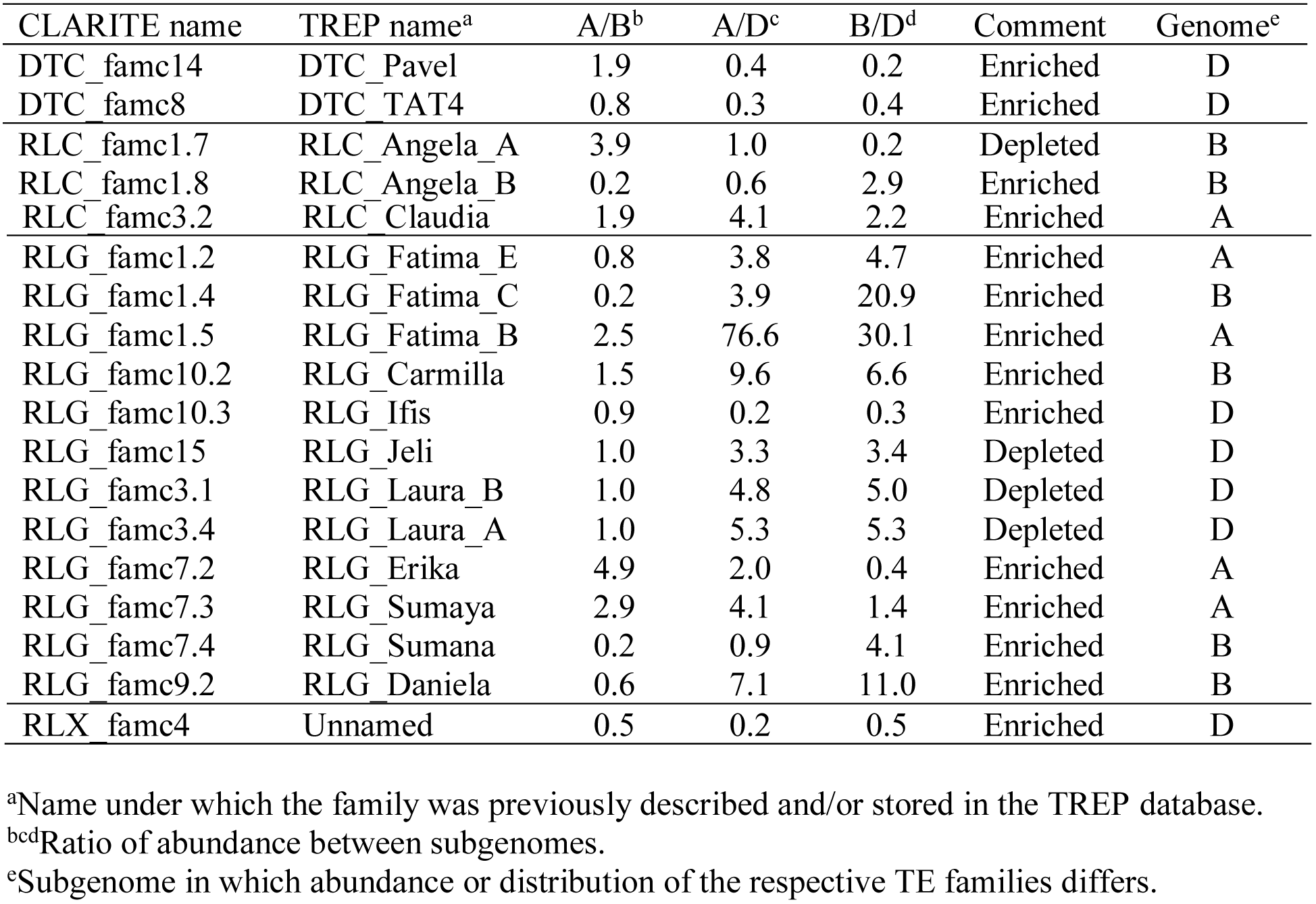
TE subfamilies that show differences in abundance between subgenomes.

### Dynamics of LTR-retrotransposons from the diploid ancestors to the hexaploid

The largest portion of plant genomes consists of long terminal-repeat retrotransposons (LTR-RTs). Intact full-length elements represent recently inserted copies whereas old elements have experienced truncations, nested insertions, and mutations that finally lead to degenerated sequences until they become un-recognizable. Full-length LTR-RTs (flLTR-RTs) are bordered by two LTRs that are identical at the time of insertion and subsequently diverge by random mutations, a characteristic that is used to determine the age of transposition events [13]. In previous genome assemblies, terminal repeats tended to collapse which resulted in very low numbers of correctly reconstructed flLTR-RTs (triangles in Fig. S13). We found 112,744 flLTR-RTs in RefSeq_v1.0 (Table S1, Fig. S13) which was in line with the expectations and confirmed the linear relationship between flLTR-RTs and genome size within the *Poaceae.* This is two times higher than assembled in TGAC_v1 [30], while almost no flLTR-RT were assembled in the 2014 gene-centric draft assembly [31].

We exploited this unique dataset to gain insights into the evolutionary history of hexaploid wheat from a transposon perspective. flLTR-RTs are evenly distributed among the subgenomes, with on average 8 elements per Mb (Table S1). Among them, there were two times more Copia (RLC) than Gypsy (RLG) elements, although Gypsy elements account for 2.8x more DNA. This is in agreement with the overall younger age of Copia TE. Indeed, the median insertion ages for Copia, Gypsy and RLX (unclassified LTR-RTs) are 0.95, 1.30 and 1.66 Myrs. RLXs lack a protein domain, preventing a straightforward classification into Gypsy or Copia.

The missing domains can most likely be accounted for by their older age and, thus, their higher degree of degeneration. RLX elements are probably unable to transpose on their own but the occurrence of such very recently transposed elements suggests that they are non-autonomous as described for Fatima subfamilies (Fig 3A). Between the A and B subgenomes, all flLTR-RT metrics are very similar, whereas the D subgenome stands out with younger insertions.

We analyzed the chromosomal distributions of the flLTR-RTs (Fig. S14). The whole set of elements is relatively evenly scattered along the chromosomes with high density spots in the distal gene-rich compartments. The most recent transpositions (age=0) involved 457 elements: 257 Copia, 144 Gypsy, and 56 RLXs. They are homogeneously distributed along the chromosomes (Fig. S14B), confirming previous hypotheses stating that TEs insert at the same rate all along the chromosome but are deleted faster in the terminal regions, leading to gene-rich and TE-depleted chromosome extremities [17].

The current flLTR-RT content is the outcome of two opposing forces: insertion and removal. Therefore, we calculated a persistence rate, giving the number of elements per 10,000 years that have remained intact over time, for the 112,744 flLTR-RTs (Fig 4A). It revealed broad peaks for each superfamily, with maxima ranging from 0.6 Mya (for Copia in the D subgenome) to 1.5 Mya (for RLX in A and B subgenomes). The D subgenome contained on average younger flLTR-RTs compared to A and B, with a shift of activity by 0.5 Myrs. Such peaks of age distributions are commonly interpreted in the literature as transposon amplification bursts. We find the “burst” analogy misleading because the actual values are very low. For wheat, it represents a maximal rate of only 600 copies per 10,000 years. A more suiting analogy would be the formation of mountain ranges where small net increases over very long-time periods add up to very large systems. In the most recent time (<10,000 years), after the hexaploidization event, we did not see any evidence in our data for the popular “genomic shock” hypothesis, postulating immediate drastic increases of transposon insertions [32-34]. For the A and B subgenomes, a shoulder in the persistence curves around 0.5 Mya (Fig. 4A), the time point of tetraploidization, was observed. We suggest that counter-selection of harmful TE insertions was relaxed in the tetraploid genome, i.e. the polyploid could tolerate insertions which otherwise would have been removed by selection in a diploid.

**Figure 4:**
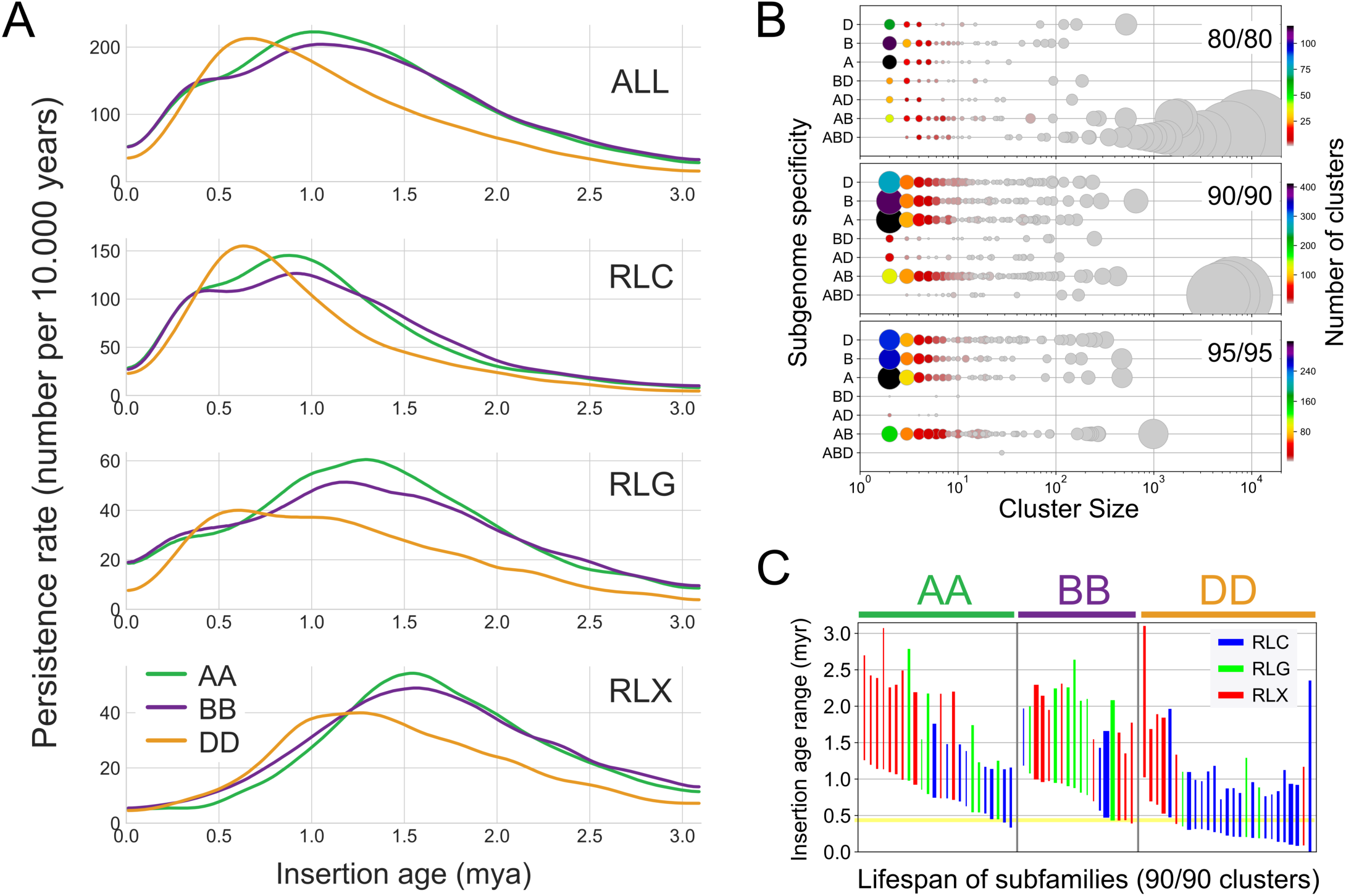
Insertion time-frames of wheat LTR-retrotransposons. **A.** Persistence rate in number of elements per 10,000 years that have remained intact until now (meaning they have not been removed or truncated over time). The D genome has younger flLTR-RTs, the curves for all superfamilies are shifted by ~0.5 Myrs. The shoulder at 0.5 Myrs in the A and B subgenomes could reflect a decrease in removal rates after the tretraploidization. **B.** Comparison of different cluster stringencies. Y-axis: subgenome specificity of the clusters, e.g. “ABD” has members from all three subgenomes, “AB” only from A and B; X-axis: log cluster size; the color coding gives the number of clusters; the circle area corresponds to the number of elements. The family clustering at 80% identity over 80% mutual coverage generates large clusters, but has a low proportion of subgenome-specific clusters. The 90/90 subfamily level cluster set with a high number of subgenome specific clusters and three large ABD clusters was used for further analyzes. **C.** Lifespan of subfamilies containing only either A, B, or D members. The line thickness represents cluster size. Lineages unique to the A or B subgenome occur only down to ~0.5 Myrs, confirming the estimated time point for the tetraploidization. However, D subgenome-unique lineages kept on proliferating, a clear sign for a very recent hexaploidization.

To elucidate the TE amplification patterns that have occurred before and after polyploidization, we clustered the 112,744 flLTR-RTs based on their sequence identity. The family level was previously defined at 80% identity over 80% sequence coverage (80/80 clusters) [2]. We also clustered the flLTR-RTs using a more stringent cutoff of 90/90 and 95/95 to enable classification at the subfamily level (Fig. 4B). 80/80 clusters were large and contained members of all three subgenomes. In contrast, the 90/90 and 95/95 clusters were smaller and a higher proportion of them are specific to one subgenome. To trace the polyploidization events, we defined lifespans for each individual LTR-RT subfamily as the interval between the oldest and youngest insertion (Fig. 4C). Subfamilies specific to either the A or B subgenome amplified until about 0.4 Myrs, which is consistent with the estimated time of the tetraploidization. Some of the D subgenome-specific subfamilies inserted more recently, again consistent with the very recent hexaploidization.

These results confirmed that the three subgenomes were shaped by common families present in the A-B-D common ancestor that have amplified independently in the diploid lineages. They evolved to give birth to different subfamilies that, generally, did not massively exchange since polyploidization. To confirm this hypothesis, we explored the phylogenetic trees of the three largest 90/90 clusters color coded by subgenome (Fig. 5 and Fig. S15-S17 for more details). The trees show older subgenome-specific TE lineages which have proliferated in the diploid ancestors (2-0.5 Mya). However, the youngest elements (<0.5 Mya) were found in clades interweaving elements of the A and B subgenomes, corresponding to amplifications in the tetraploid. Such cases involving the D subgenome were not observed, showing that flLTR-RTs from D have not yet transposed in large amounts across the subgenomes since the birth of hexaploid wheat 8000-10000 years ago. We further noticed several incidences in the trees where D lineages were derived from older B or A lineages, but not the reverse. This may be explained by the origin of the D subgenome through homoploid hybridization between A and B [35].

**Figure 5:**
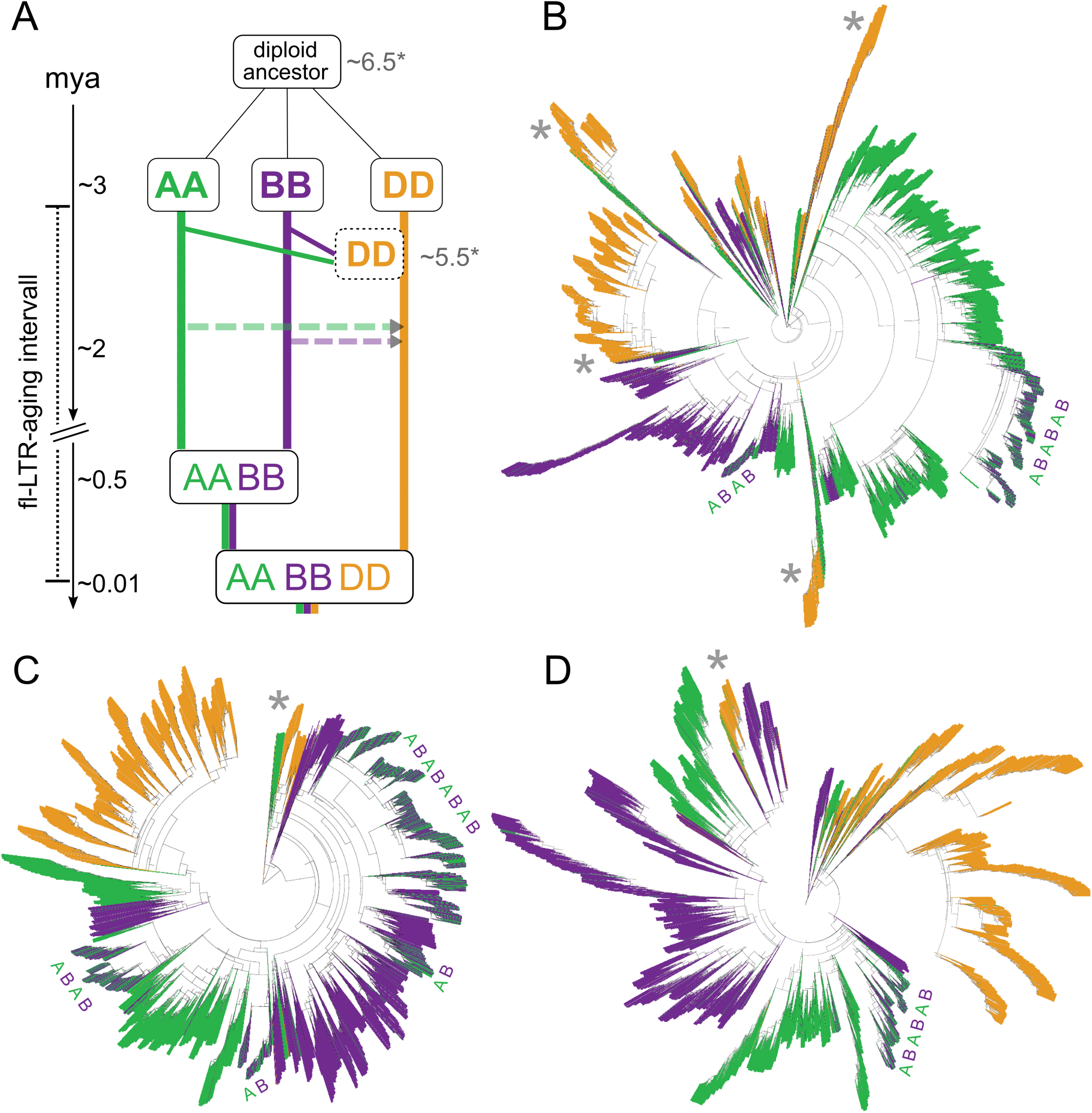
LTR-retrotransposon footprints in the evolution of hexaploid wheat. **A.** Evolution of the wheat genome with alternative scenarios and timescales. The dotted rectangles and ^∗^ time values represent the scenario of A and B giving rise to the D subgenome by homoploid hybridization [35]. The left timescale is based on another estimate based on the chloroplast genome evolution [16]. The dotted horizontal arrows represent the unidirectional horizontal transposon transfers observed in this study. **B**. Phylogenetic tree of the largest 90/90 cluster (6,639 copies). **C.** Top2 cluster (5,387 copies), **D.** Top3 cluster (4,564 copies). The leaves of the tree are colored by the subgenome localization of the respective elements. The majority of the amplifications took place in the diploid ancestors evidenced by the single colored propagation lineages. Each tree contains one or several younger regions with interweaving A and B insertions (marked by ABAB). These younger proliferations only started in the AABB tetraploid, where the new elements inserted likewise into both subgenomes. The joining of the D genome was too recent to have left similar traces yet. The gray asterisks mark D lineages that stem from a B or A lineage.

There are two proposed models of propagation of TEs the “master copy” and “transposon” model [36]. The “master copy” model gives rise to highly unbalanced trees (i.e. with long successive row patterns) where one active copy is serially replaced by another, whereas the “transposon” model produces balanced trees where all branches duplicate with the same rate [37]. To better discern the tree topologies, we plotted trees with equal branch length and revealed that the three largest trees (comprising 15% of flLTR-RTs) are highly unbalanced (Fig. S18) while the smaller trees are either balanced or unbalanced (Fig. S19). Taken together, both types of tree topologies exist in the proliferation of flLTR-RTs, but there is a bias towards unbalanced trees for younger elements, suggesting that TEs proliferation followed the “master copy” model.

In summary, our findings give a timed TE atlas depicting detailed TE proliferation patterns of hexaploid wheat. They also show that polyploidization did not trigger bursts of TE activity. This dataset of well-defined transposon lineages now provides the basis to further explore the genetic and regulatory aspects of fine scaled transposon dynamics and to answer intriguing questions like: which factors cause a lineage to start and what causes activity to stop?

### A stable genome structure despite the massive reshuffling of intergenic sequences

Despite the massive turnover of the TE space (Fig. 2A), the gene order along the homeologous chromosomes is well conserved between the subgenomes and is even conserved with the related grass genomes (sharing a common ancestor 60 Mya [38]). Most interestingly and strikingly, not only gene order but also distances between neighboring homeologs tend to be conserved between subgenomes (Fig. 6). Indeed, we found that the ratio of distances between neighboring homeologs has a strong peak at 1 (or 0 in log scale on Fig. 6), meaning that distances separating genes tend to be conserved between the three subgenomes despite the TE turnover. This effect is non-random, as ratio distribution curves are significantly flatter when gene positions along chromosomes are randomized. These findings suggest that the overall chromatin structure is likely under selection pressure.

**Figure 6:**
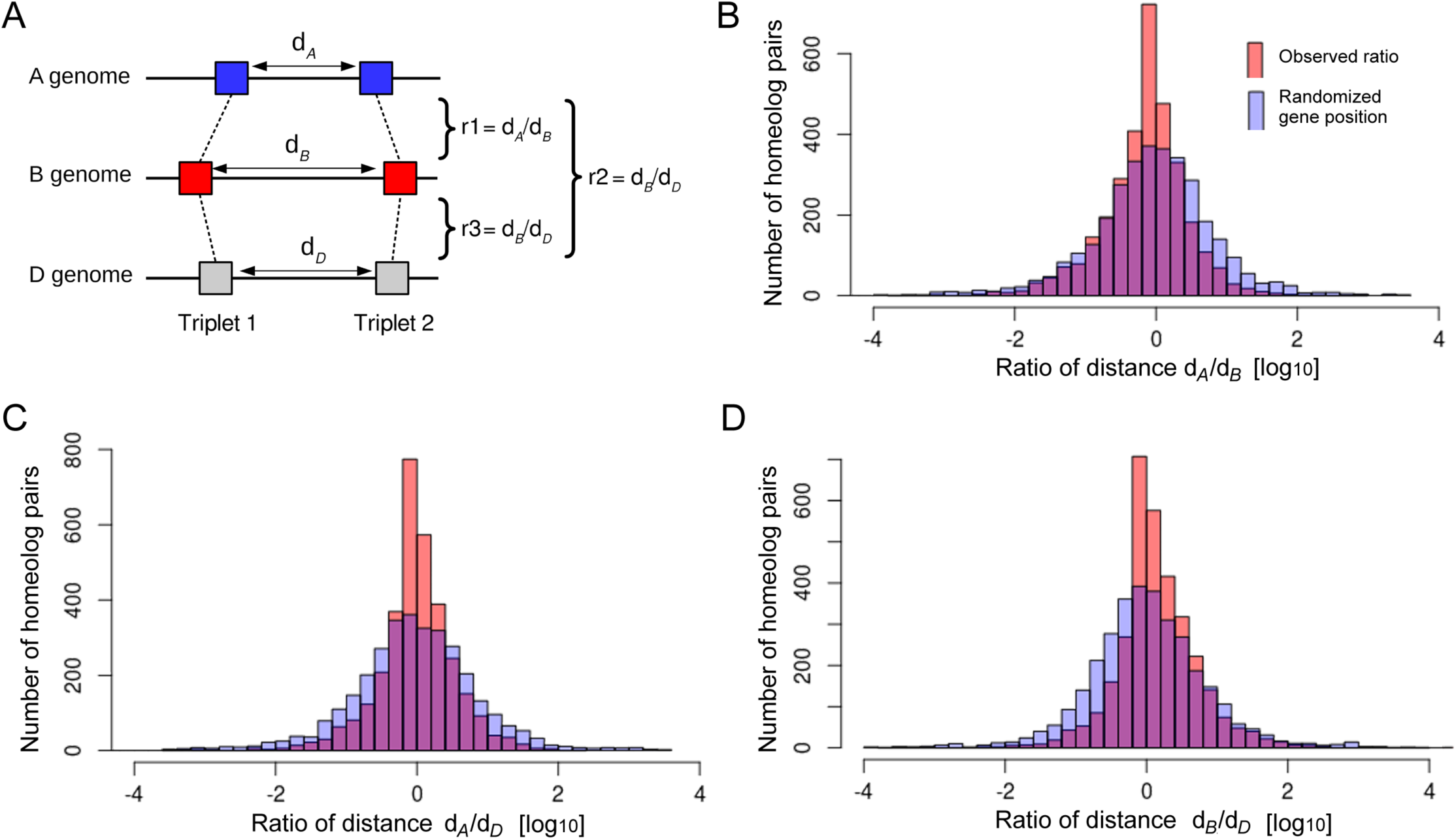
Comparison of distances between neighboring homeologs in the subgenomes. **A.** Distances between genes and their closest neighbors were compared to those of their homeologous partners from the other subgenomes. For each homeolog triplet, three ratios were calculated (i.e. pairwise comparisons between the three subgenome homeologs). If the distance is similar in two subgenomes, the ratio will be close to 1. **B**. Comparison of 2,275 gene pairs from the terminal 150 Mb of short chromosome arms from A and B genomes. The distribution is compared to one where gene positions were randomized (see methods). The observed data has a sharper peak at 1 (logarithmic scale where log(1)=0). This indicates that distances between homeologs are conserved, despite the near-complete absence of conservation of intergenic sequences between subgenomes. **C**. Analogous comparison of homeolog pairs from the A and D genomes. **D**. Analogous comparison of homeolog pairs from the B and D genomes.

We found this constrained distribution irrespective of the chromosome compartments i.e. distal, interstitial, and proximal, exhibiting contrasted features at the structural (gene density) and functional (recombination rate, gene expression breadth) levels [24, 25]. However, constraints applied on intergenic distances seem relaxed (broader peak in Fig. 6) in proximal regions where meiotic recombination rate is extremely low. At this point, we can only speculate about the possible impact of meiotic recombination as a driving force towards maintaining a stable chromosome organization. Previous studies have shown that recombination in highly repetitive genomes occurs mainly in or near genes [39]. Thus, it might be important that spacing of genes is preserved for proper expression regulation or proper pairing during meiosis. Previous studies on introgressions of divergent haplotypes in large-genome grasses support this hypothesis. For instance, highly divergent haplotypes which still preserve the spacing of genes have been maintained in wheats of different ploidy levels at the wheat *Lr10* locus [40].

### Presence of genes had a major impact on the TE content, and a similar impact between the A, B, and D subgenomes

The sequences directly surrounding genes have a distinct TE composition compared to the overall TE space. While large intergenic regions are dominated by large TEs such as LTR-RTs and CACTAs, sequences surrounding genes are enriched in small TEs that are usually just a few hundred bps in size (Fig. 7). Even in the narrow sequence window of 2 kb up- and downstream of genes (i.e. promoters and regulatory regions of protein-coding genes), we found sub-compartments which are defined by the enrichment of specific superfamilies. Immediately up- and downstream of genes, we identified mostly small non-autonomous DNA transposons of the Mariner superfamily. These small TEs are also referred to as Stowaway MITEs [41]. Slightly further upstream, peaks of SINEs and Mutator were also clearly visible (Fig. 7). Approximately, 1.4 kb upstream of predicted transcription start sites (TSS), Harbinger elements have their highest enrichment. These small non-autonomous elements have been referred to as Tourist MITEs [41]. Interestingly, downstream regions differ in their composition from upstream regions. Here, SINEs, Mutator and Mariner elements are clearly less abundant, which makes the TE landscape surrounding genes asymmetrical (Fig. 7). Again, although the TE content of A-B-D subgenomes evolved independently, the TE landscape in the direct vicinity of genes is very similar between the three subgenomes at the superfamily level (Fig. 7, Fig. S20).

**Figure 7:**
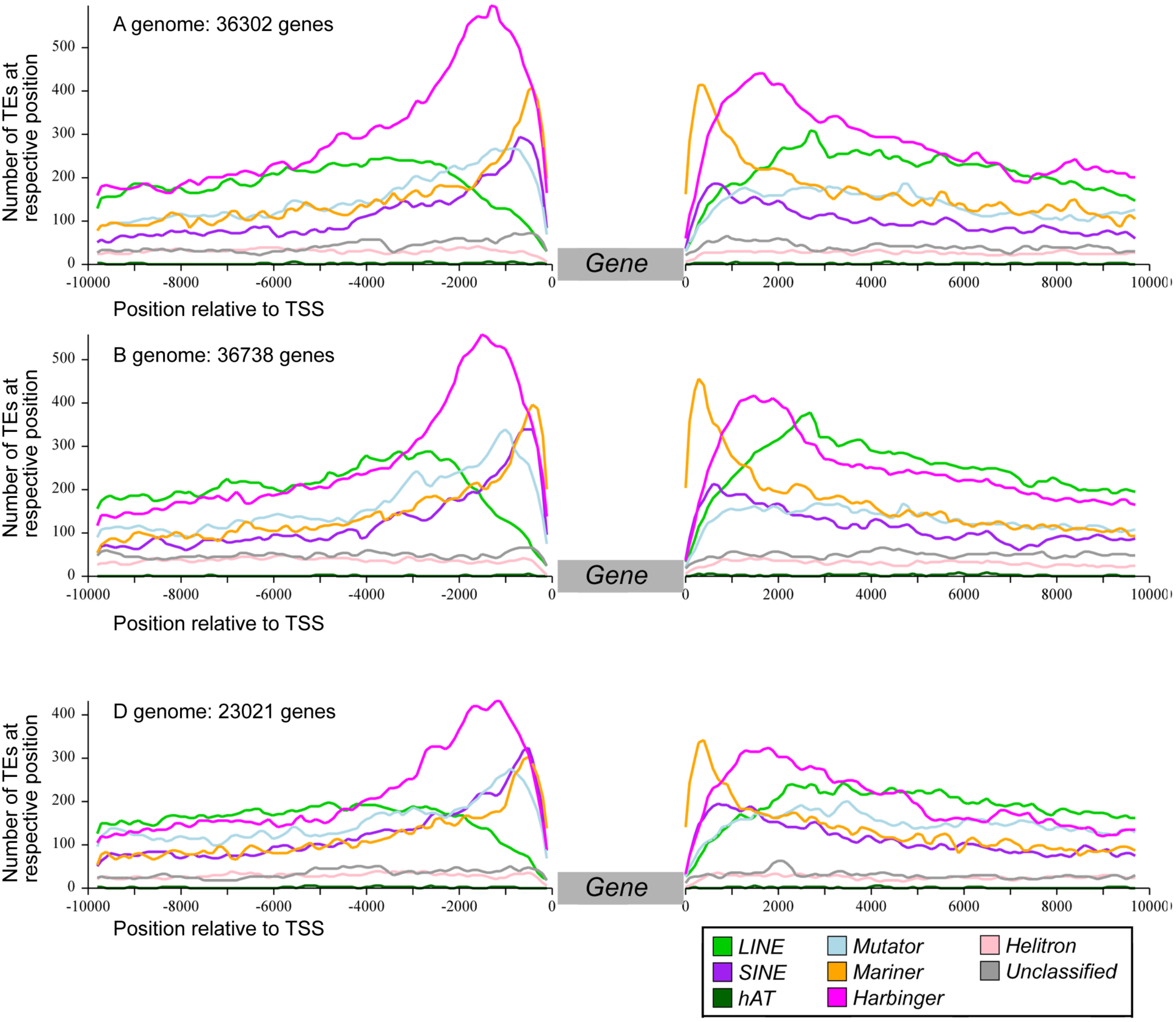
TE landscape surrounding genes. Genes from the three subgenomes were treated separately. For all genes, the 10 kb upstream of the transcription start site (TSS) and 10 kb downstream of the transcription end site were analyzed. Abundance of the different TE families was compiled for all genes of each subgenome. The plots include only those superfamilies that are specifically enriched near genes and which are otherwise less abundant in intergenic sequences.

We then computed, independently for each subgenome, the enrichment ratio of all TE families that were present in the promoter of protein-coding genes (2 kb upstream to the TSS) compared to their overall proportion (in copy number). The purpose was to identify how many and which families are over- or under-represented in gene promoters and if such preferential insertion pattern is similar in A, B, and D. To avoid biases due to small samples, we only considered the 315 families with at least 500 repeat copies at the whole genome level. The vast majority (242, 77%) showed a bias (i.e. at least a 2-fold difference in abundance) in gene promoters compared to their subgenome average. This clear difference showed that the direct physical environment of genes contrasts from the intergenic space, likely causing two distinct genomic compartments reminiscent of the distinction between euchromatin and heterochomatin. We focused on families for which the bias is strong i.e. at least 3-fold over- or underrepresentation in promoters compared to the subgenome average. In total, 105 (33%) and 38 (12%) families, respectively, met this threshold in at least one subgenome. Thus, for almost half of the TE families, their presence along the genome is driven either positively or negatively by the presence of genes. Most strikingly, this significant bias was found for all three subgenomes. While it was previously known that MITEs (namely small non-autonomous elements of the Mariner and Harbinger superfamilies) were enriched in promoters of genes, here we show that such bias is not restricted to MITEs but rather involves TEs of several different superfamilies. These results suggest that the chromatin architecture is under selection leading to a highly organized genome in which each TE family has its own genomic niche.

Again, the promoter-prone *versus* promoter-excluded pattern was extremely conserved between A, B, and D subgenomes (Fig. 8) although individual TE insertions are not conserved between homeologous promoters. In other words, when a family is over- or under-represented in the promoter regions of one subgenome, it is also true for the two other subgenomes, meaning that the specificity of insertion in genic or non-genic regions is an intrinsic ancestral feature for each family, that has been maintained independently in the three lineages. We did not find any family that was enriched in gene promoter in a subgenome while under-represented in gene promoters of another subgenome (even at a 2-fold change threshold).

**Figure 8:**
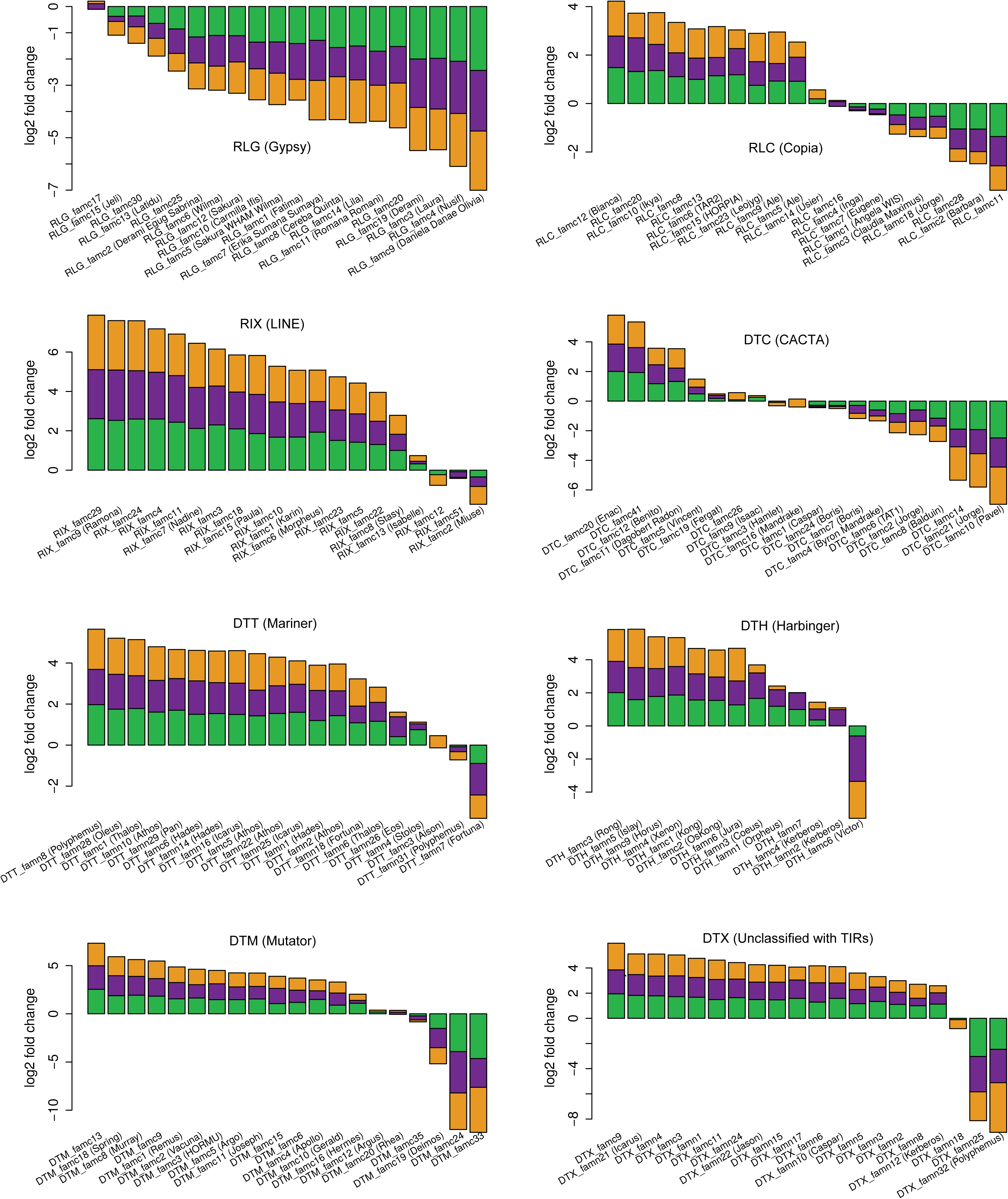
Enrichment analyses of TE families within gene promoters. Y axis represents the log2ratio of the proportion (i.e. percentage in term of number of copies) of each TE family observed in the promoter of genes (2 kb upstream the TSS) relatively to their proportion at the whole subgenome level. Positive and negative values represent an over- and under-representation of a given family in the promoters, respectively. Log2ratio were calculated for the three subgenomes independently (A: green; B: violet; D: orange) and the 3 values were represented here as a stacked histogram. Only highly repeated families (500 copies or more) are represented, with 1 panel per superfamily. Families are ordered decreasingly along the x axis according to the whole genome log2ratio.

TE families over- or under-represented in gene promoters belong to almost all superfamilies (Fig. 8), meaning that each family has its own behavior which is not necessarily conserved at the superfamily level. However, superfamily is generally a good indicator of the presence/absence of TEs in genic regions. For instance, 83% (25/30) of the LINE families are overrepresented in the promoter regions while none of them are underrepresented (considering 2fold change). Enrichment analysis also confirmed the strong tendency previously suggested that class 2 DNA transposons (especially MITEs) are enriched in promoters while Gypsy retrotransposons tend to be the main contributors of the intergenic space. Indeed, among the 105 families strongly enriched in promoters (3-fold change), 53% (56) are from class 2 and 21% (22) are LINEs, and only 5% (5) are a LTR-retrotransposons. In contrast, among the 38 families underrepresented in promoters, we found mainly Gypsy families. Figure 8 presents the fold change enrichment relatively to gene promoters. Contrary to Gypsy, Mutator, Mariner, and Harbinger, families belonging to CACTA and Copia superfamilies do not share a common specificity of insertion. We found similar proportions of them that can be either over- or underrepresented in promoters. This confirmed previous results about CACTAs annotated along the 3B chromosome [17] revealing that a part of the CACTA families is associated with genes while the other follows the distribution of Gypsy. Our results showed that it is the same for the Copia.

Thus, almost all TE families tend to be at the equilibrium in the genome, with amplification compensating their deletion, and, moreover, each family has conserved its intrinsic affinity with genes. To explain such unexpected results, we suggest that TEs would play a role at the structural level to maintain an intact chromosome architecture. This is well known for the centromeres and telomeres and we suggest that, for complex genomes, this is also true for the rest of the genome.

### No strong association between gene-flanking TEs and gene expression pattern

We investigated the influence of neighboring TEs on gene expression. Indeed, TEs are so abundant in the wheat genome, that genes are almost systematically flanked by a TE in the direct vicinity. The median distance between the gene transcription start site (TSS) and the closest upstream TE is 1.52 kb and the median distance between the transcription termination site (TTS) and the closest downstream TE is 1.55 kb, while the average gene length (between TSS and TTS) is 3.44 kb. The density as well as the diversity of TEs in the vicinity of genes allows to speculate on potential relationships between TEs and gene expression regulation. We used the gene expression network built by [25] based on an exhaustive set of wheat RNA-Seq data. Genes were clustered into 39 expression modules sharing a common expression profile across all samples.

We also grouped unexpressed genes to study the potential influence of TEs on neighbor gene silencing. For each gene, the closest TE upstream was retrieved and we investigated potential correlations though an enrichment analysis (each module was compared to the full gene set). Despite the close association between genes and TEs, no strong enrichment for a specific family was observed for any module, neither for the unexpressed genes.

We then studied the TE landscape upstream of wheat homeolog triplets, focusing on 19,393 triplets (58,179 genes) with a 1:1:1 orthologous relationship between A, B, and D subgenomes. For each triplet, we retrieved the closest TE flanking the TSS and investigated the level of conservation of flanking TEs between homeologs. For 75% of the triplets, the three flanking TEs belong to three different families, revealing that, even in the close vicinity of genes, TEs are in majority not conserved between homeologs due to rapid turn-over. This suggests that most TEs present upstream of triplets were not selected for by the presence of common regulatory elements across homeologs. However, for 736 triplets (4%), the three homeologs are flanked by the same element, constituting a conserved noncoding sequence (CNS), suggesting that part of this element is involved in the regulation of gene expression. These TE-derived CNS are on average 459 bp which is 3 times smaller than the average size of gene-flanking TE fragments (on average 1,355 bp), suggesting that only a portion of the ancestrally inserted TE is under selection pressure. They represent a wide range (149 different families) of diverse elements belonging to all the different superfamilies.

Given that the majority of homeolog triplets have relatively similar expression patterns [25, 42] we next focused on triplets which had distinct expression profiles. We examined transcriptionally dynamic triplets [42] across tissues and triplets for which two copies are coexpressed while the third is not expressed. For both datasets, enrichment analysis did not reveal any significant enrichment between expression profiles and the presence of a particular TE family in the promoter. These results suggest that the expression pattern of genes in wheat is not determined by the TEs present in the close vicinity of the genes.

## Conclusions

The chromosome-scale assembly of the wheat genome provided an unprecedented genome-wide view of the organization and impact of TEs in such a complex genome. Since they diverged, the A, B, and D subgenomes have experienced a near-complete TE turnover although polyploidization did not massively reactivate TEs. This turnover contrasted drastically with the high level of gene synteny. Apart from genes, there was no conservation of the TE space between homeologous loci. But surprisingly, TE families that have shaped the A, B, and D subgenomes are the same, and unexpectedly, their proportions and intrinsic properties (gene-prone or not) are quite similar despite their independent evolution in the diploid lineages. Thus, TE families are somehow at the equilibrium in the genome since the A-B-D common ancestor. These novel insights contradict previous model of evolution with amplification bursts followed by rapid silencing. Our results suggest a role of TEs at the structural level with a chromatin architecture under selection pressure. TEs are not just junk DNA and our findings open new perspectives to elucidate their role on high-order chromatin arrangement, chromosome territories, and gene regulation.

## Methods

### TE modeling using CLARITE

The *Triticum aestivum* cv. Chinese Spring genome sequence was annotated as described in [25]. Briefly, two gene prediction pipelines were used (TriAnnot: developed at GDEC Institute [INRA-UCA Clermont-Ferrand]; the pipeline developed at Helmholtz Center Munich [PGSB]) and the two annotations were integrated (pipeline established at Earlham Institute [43]) to achieve a single high-quality gene set. TE modeling was achieved through a similarity search approach based on the ClariTeRep curated databank of repeated elements [44], developed specifically for the wheat genome, and with the CLARITE program that was developed to model TEs and reconstruct their nested structure [17]. For the annotation, we used the ClariTeRep naming system which assigns simple numbers to individual families and subfamilies e.g. RLG_famc1.1 and RLG_famc1.2 are subfamilies of RLG_famc1. Since many TE families have been previously named, we provided this name in parentheses.

### Detection and characterization of full-length LTR-retrotransposons

Identification of flLTR-RTs was based on LTRharvest [45]. For the RefSeq_v1.0, LTRharvest reported 501,358 non-overlapping flLTR-RT candidates under the following parameter settings: “overlaps best -seed 30 -minlenltr 100 -maxlenltr 2000 -mindistltr 3000 - maxdistltr 25000 -similar 85 -mintsd 4 -maxtsd 20 -motif tgca -motifmis 1 -vic 60 -xdrop 5 -mat 2 -mis −2 -ins −3 -del −3”. All candidates where annotated for PfamA domains with hmmer3 [46] and stringently filtered for canonical elements by the following criteria: i. presence of at least one typical retrotransposon domain (RT, RH, INT, GAG); ii. removal of mis-predictions based on inconsistent domains e.g. RT-RH-INT-RT-RH; iii. absence of gene-related Pfam domains; iv. strand consistency between domains and primer binding site; v. tandem repeat content below 25%; vi. long terminal repeat size <=25% of the element size; vii. N content <5%. This resulted in a final set of 112,744 high quality flLTR-RTs. The Copia and Gypsy superfamilies where defined by their internal domain ordering: INT-RT-RH for RLC and RH-RT-INT for RLG [2]. When this was not possible, the prediction was classified as RLX. The 112,744 flLTR-RTs were clustered with vmatch dbcluster [47] at three different stringencies: 95/95 (95% identity over 95% mutual length coverage), 90/90, and 80/80, as follows: vmatch “-dbcluster 95 95 -identity 95 -exdrop 3 -seedlength 20 -d”, “-dbcluster 90 90 -identity 90 -exdrop 4 -seedlength 20 -d” and “-dbcluster 80 80 -identity 80 -exdrop 5 -seedlength 15 -d”. Subgenome specificity of clusters was defined by the following decision tree: i. assignment of the respective subgenome if >=90% of the members where located on this subgenome; ii. assignment to two subgenomes if members from one subgenome <10%, e.g. AB-specific if D members <10%; iii. assignment of the remaining clusters as ABD common. Muscle was used for multiple alignments of each cluster [48] in a fast mode (-maxiters 2 -diags1). To build phylogenetic trees, we used tree2 from the muscle output which was created in the second iteration with a Kimura distance matrix and trees were visualized with ete3 toolkit [49]. The date of flLTR-RT insertions was based on the divergence between the 5’ and 3’ LTRs calculated with emboss distmat, applying the Kimura 2-parameter correction. The age was estimated using the formula: age=distance/(2^∗^mutation rate) with a mutation rate of 1.3*10-8 [13]. The lifespan of an individual LTR-RT subfamily was defined as the 5 to 95 percentile interval between the oldest and youngest insertions. The densities for the chromosomal heat-maps where calculated using a sliding window of 4 Mb with a step of 0.8 Mb.

### Comparative analysis of distances separating neighbor genes between homeologous chromosomes

For the comparison of distances separating neighbor genes, homeologous triplets located in the three chromosomal compartments (distal, interstitial, and proximal; Table S2) were treated separately. This was done because gene density is lower in interstitial and proximal regions, and because these latter show a lack of genetic recombination. Furthermore, we considered only triplets where all three homeologous genes are found on the homeologous chromosomes. Comparison of homeologous gene pairs from distal regions was done in two ways which both yielded virtually identical results. Distances were measured from one gene to the one that follows downstream. However, there were many small local inversions between the different subgenomes. Thus, if a gene on the B or D subgenomes was orientated in the opposite direction compared to its homeologous copy in the A subgenome, it was assumed that that gene is part of a local inversion. Therefore, the distance to the preceding on the chromosome was calculated. The second approach was more stringent, based only on triplets for which all three homeologs are in the same orientation in the three subgenomes. The results obtained from the two approaches were extremely similar and we presented only the results from the second, more stringent, approach. For the control dataset, we picked a number of random positions along the chromosomes that is equal to the number of homeologs for that chromosome group. Then, homeologous gene identifiers were assigned to these positions from top to bottom (to preserve the order of genes but randomize the distances between them). This was done once for all three chromosomal compartments.

### Analyses of TEs in the vicinity of genes and enrichment analyses

We developed a Perl script (gffGetClosestTe.pl) to retrieve gene-flanking TEs from the feature coordinates in GFF file. It was used to extract the closest TE on each side of every predicted gene (considering “gene” features that includes UTRs). It was also used to extract all predicted TE copies entirely or partially present within 2 kb upstream the “gene” start position i.e. the transcription start site (TSS). Enrichment analyses were then automated using R scripts. *1. Enrichment of TE families in gene promoters (2 kb upstream).* Independently for the three subgenomes, we retrieved all TE copies present with 2 kb upstream the TSSs of all gene models and calculated the percentage of the number of copies assigned to every family (%famX^promoter^). We also calculated the percentage of the number of copies of each family at the whole subgenome level (%famX^whole_subgenome^). One enrichment log2ratio was calculated for each A, B, and D subgenome using the formula: log2(%famX^promote^7%famX^wbole_subgenome^). Only families accounting for 500 copies or more in the whole genome were considered. *2. TE families and expression modules.* Here, we retrieved the closest TE present in 5’ of the TSS for all genes and calculated the percentage of each TE family for each expression module and the unexpressed genes (considered as a module), and compared them to the percentage observed for the whole gene set using the formula: log2(%famX^genes^-^moduleX^/%famX^all_genes^). Log2ratio was calculated only for expression modules representing at least 1,000 coexpressed genes and we considered only log2ratio values for families accounting for 500 copies or more. A similar approach was taken for the 10% stable, 80% middle and 10% dynamic genes as defined by [42]. *3. Comparison of TE families in the promoter of homeologs.* Here, we also retrieved the closest TE in 5’ of every gene and identified homeologous triplets for which the closest element in 5’ belongs to the same family for the three copies. For that, we developed a Perl script (getTeHomeologs.pl) in order to integrate the information of homeologous genes and the (closest TE in 5’ of genes. Only “1-1-1” homeologs were considered.

## List of abbreviations

CNS: conserved non-coding sequence
flLTR-RT: full-length long terminal repeat retrotransposon
INT: integrase
LINE: long interspersed nuclear element
LTR: long terminal repeat
MITE: miniature inverted-repeat transposable element
ORF: open reading frame
RH: ribonuclease H
RT: retrotransposon
SINE: short interspersed nuclear element
TE: transposable element
TSS: transcription start site
TTS: transcription termination site

## Declarations

### Ethics approval and consent to participate

Not applicable

### Consent for publication

Not applicable

### Availability of data and material

Data availability: IWGSC RefSeq_v1.0 assembly, gene and TE annotation as well as related data are available on the IWGSC Data Repository hosted at URGI: https://wheat-urgi.versailles.inra.fr/Seq-Repository.

## Authors’ contributions

TW, HG, and FC conceived and designed the study. FC coordinated the study. TW performed analyses of TE distribution along chromosomes, comparisons of intergenic distances between subgenomes, centromeric TE analyses, and analyzed the TE landscape around genes. HG carried out the kmer analyses, flLTR-RT identification, dating, clustering and phylogenetic analyses. FC carried out the A-B-D comparisons of TE families, enrichment analyses in gene promoters, comparisons between homeologs, and association with expression profiles. CU, RHRG, and PB built the RNA-Seq based gene expression network. TW, HG and FC wrote the manuscript. RDO contributed text and figures for the manuscript. MS, CU, RHRG, PB, KFXM, RDO, and EP provided comments and corrections on the manuscript.

## Acknowledgements

Not applicable

## Additional Files

**Additional File 1**: Supplementary Tables S1 to S2, Figures S1 to S20

